# Protein phase separation provides long-term memory of transient spatial stimuli

**DOI:** 10.1101/283804

**Authors:** Elliot Dine, Agnieszka A. Gil, Giselle Uribe, Clifford P. Brangwynne, Jared E. Toettcher

## Abstract

Protein/RNA clusters arise frequently in spatially-regulated biological processes, from the asymmetric distribution of P granules and PAR proteins in developing embryos to localized receptor oligomers in migratory cells. This co-occurrence suggests that protein clusters might possess intrinsic properties that make them a useful substrate for spatial regulation. Here, we demonstrate that protein droplets show a robust form of spatial memory, maintaining the spatial pattern of an inhibitor of droplet formation long after it has been removed. Despite this persistence, droplets can be highly dynamic, continuously exchanging monomers with the diffuse phase. We investigate the principles of biophysical spatial memory in three contexts: a computational model of phase separation; a novel optogenetic system where light can drive rapid, localized dissociation of liquid-like protein droplets; and membrane-localized signal transduction from clusters of receptor tyrosine kinases. Our results suggest that the persistent polarization underlying many cellular and developmental processes could arise through a simple biophysical process, without any additional requirement for biochemical positive and negative feedback loops.

**Highlights:**

1. We introduce PixELLs, an optogenetic system for protein droplet disassembly.
2. Modeling and experiments demonstrate long-term memory of local droplet dissociation.
3. Droplets ‘remember’ spatial stimuli in nuclei, the cytosol and on cell membranes.
4. FGFR-optoDroplets convert transient local inputs to persistent cytoskeletal responses.

## Introduction

Across many biological contexts, cells must be able to sense external spatial cues and generate asymmetric distributions of their internal components. Anisotropic patterns of protein/RNA localization play crucial roles during embryo development (Kloc and Etkin, 2005; Sailer et al., 2015), and motile cells can migrate by generating persistent internal asymmetries even in a uniform environment (Prentice-Mott et al., 2016). It is often assumed that both the establishment and maintenance of these persistent spatial patterns require complex genetic and/or biochemical networks, such as Turing-like mechanisms that combine short-range positive feedback with long-range negative feedback (Gierer and Meinhardt, 1972; Turing, 1990) or stochastic processes that rely on depleting a limiting pool of proteins that participate in an auto-regulatory positive feedback loop (Altschuler et al., 2008).

Intriguingly, many spatially-regulated biological processes also exhibit hallmarks of protein phase separation, a process where multivalent interactions between monomers drive large-scale assembly into liquid-like droplets or solid aggregates (Figure 1A). Developmental processes rely on localized RNA and/or protein aggregation, including the asymmetric partitioning of PAR proteins (Goldstein and Macara, 2007), RNA granules in *Drosophila* embryogenesis (Forrest and Gavis, 2003), and P granules that dissolve and condense along the anterior-posterior axis of *C*. *elegans* embryos to be inherited by cells that form the germline (Brangwynne et al., 2009). Similar principles may also underlie spatially-restricted signaling in differentiated cells. Localized clustering of membrane receptors is thought to promote actin nucleation during cell migration (Banjade and Rosen, 2014), and local clustering of signaling proteins was recently shown to enhance signaling downstream of T cell receptor activation at the immunological synapse (Su et al., 2016). In many of the above examples, clustering is primarily thought of as playing a biochemical role: segregating proteins away from undesired interaction partners or increasing reaction rates between components that are co-localized within the separated phase (Shin and Brangwynne, 2017).

**Figure 1.**
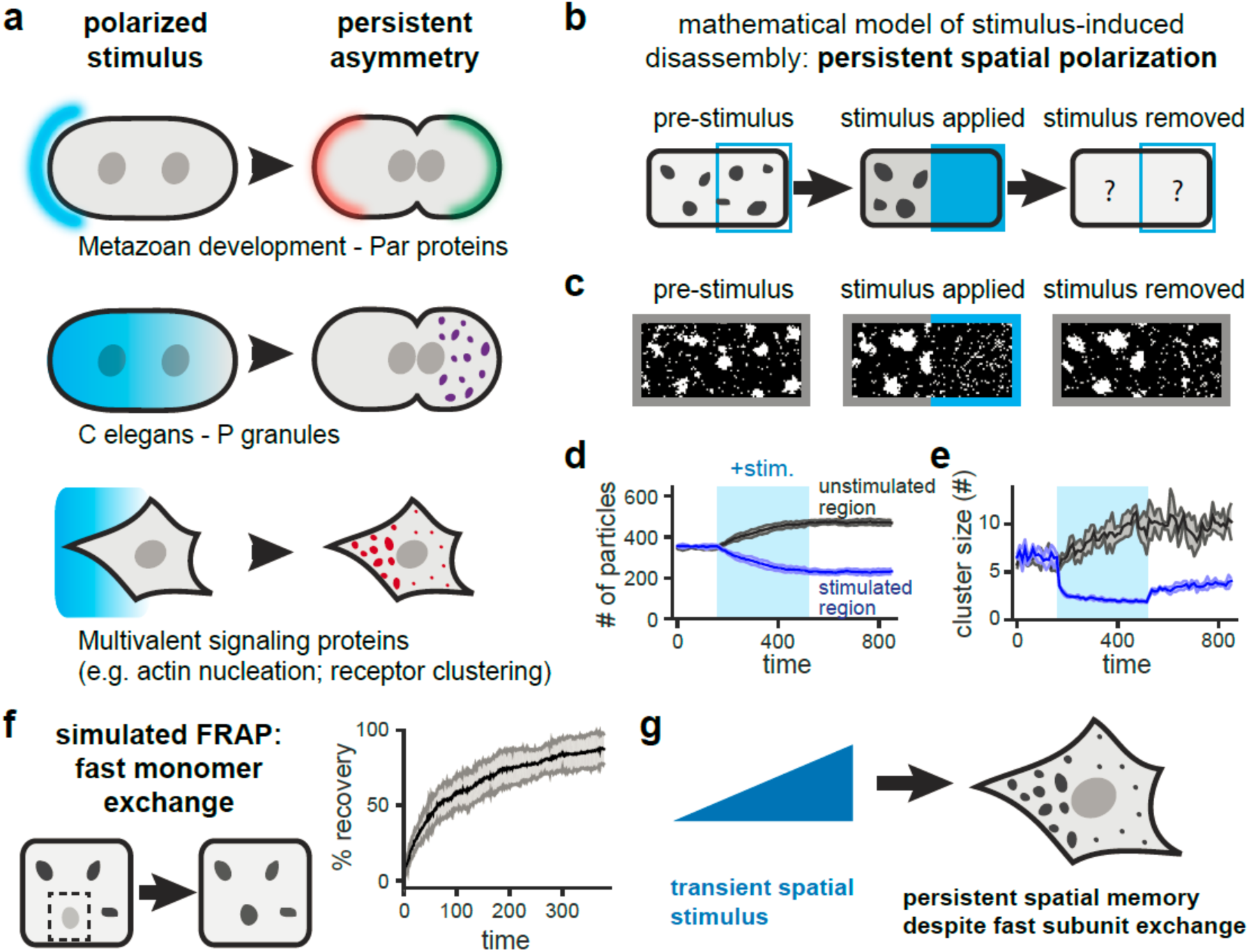
A mathematical model predicts long-term spatial memory from phase separation. (**a**) Asymmetric protein clustering occurs as part of polarized intracellular processes. (**b**) Schematic of simulated experiment where clusters are locally dissolved by a transient stimulus. (**c**) Still frames from simulation demonstrating the response to the stimulus in **b**. (**d**,**e**) Quantification of the total number of particles (**d**) and mean cluster size (**e**) in the stimulated and unstimulated regions during all three stimulation time periods. Mean + SEM are shown from five independent runs. (**f**) A simulated photobleaching experiment demonstrates rapid exchange of monomers in and out of clusters. Mean + SEM for 10 clusters is shown. (**g**) Modeling suggests that transient, local stimuli can drive persistent asymmetries of dynamic, liquid-like granules.

Here, we set out to investigate whether protein phase separation might directly contribute to the establishment or maintenance of spatial patterns within the cell. Using a combination of mathematical modeling and optogenetic stimulus experiments, we found that liquid-like protein droplets exhibit a form of long-term spatial memory. A cluster-dissociating stimulus that is delivered on one side of a cell can drive asymmetric patterns of protein localization in minutes, and these patterns persist for hours after the stimulus is removed. These results hold even in cases where droplets are highly dynamic and exchange substituents with the surrounding diffuse phase. We find that this spatial memory is robust, occurring in all three subcellular compartments tested (cytosol, nucleus, and plasma membrane) and with both optogenetic systems we employ. Finally, we show that this spatial memory can have functional implications using light-controllable FGF receptors whose phase separation drives a cytoskeletal response. Our results demonstrate that the biophysical phenomenon of protein clustering can function as a highly sensitive intracellular ‘memory foam’, amplifying transient, shallow gradients into sharp and persistent responses.

## Results

### A minimal model to dissect the role of clustering in spatial patterning

To gain some initial intuition about how phase separation might influence spatial patterning, we constructed a simple computational model of protein diffusion and clustering in two dimensions (Figure S1A; Supplementary Methods) (Freeman Rosenzweig et al., 2017; Landau and Binder, 2014). Our model consists of a 50 × 100 unit grid where each square can be occupied by a single “monomer” that is free to diffuse to adjacent unoccupied positions or exchange positions with a monomer in a neighboring occupied square. To model protein phase separation and aggregation, monomers occupying adjacent squares exhibit affinity for one another, leading to a decreased probability of movement to squares that require bond breakage. We used a temperature-like stimulus parameter θ to control the strength of binding; θ can be raised or lowered at any spatial position on the grid and at any time. Simulating the model for different values of θ revealed that it could reproduce classic properties of phase separation, including a single diffuse phase at high θ, coexistence of dynamic, liquid-like droplets and a diffuse phase at intermediate values of θ, and strongly arrested dynamics for low θ (Movie S1; Figure S1B-D) (Freeman Rosenzweig et al., 2017; Landau and Binder, 2014). Local stimulation could also drive local phase separation: decreasing θ on one half of the grid induced the appearance of local clusters that were quickly reversed when θ was returned to its initial value (Figure S1E-H).

Remarkably, even this simple model could generate complex behavior when subjected to certain classes of spatial stimuli. One illustrative example is the converse of the local stimulus experiment described above: starting from an initial state where droplets appear throughout the grid, we locally increased θ to induce droplet disassembly in a stimulated region (Figure 1B,C; Movie S2). This local stimulation led to the rapid dissolution of droplets in the stimulated area and nucleation/growth of droplets in the unstimulated region. Notably, after stimulus removal, the system did not return to its initial state but instead retained an asymmetric spatial distribution of clusters. This persistent asymmetry was visually striking and could be quantitatively captured in both the distribution of the total number of particles and the mean cluster size in the stimulated and unstimulated regions (Figure 1D,E). Importantly, this persistent asymmetry still arose even under liquid-like conditions where clusters are highly dynamic. Performing a computational FRAP experiment revealed that monomers were exchanged rapidly between clusters and the diffuse phase, even as the overall spatial distribution of clusters was unchanged (Figure 1F).

Our simple model thus suggests that in the case of stimulus-induced *dissociation* of protein clusters, long-term spatial patterns can result even from the application of a brief stimulus. This phenomenon arises from the well-characterized physics of droplet phase behavior (Doi, 2013). When droplets are dissolved by a local stimulus the concentration of monomers in that region rises, leading to a diffusive flux toward the unstimulated region, condensation into droplets there, and a return of the monomer concentration to near its pre-stimulus level. Upon stimulus removal, however, there is no driving force for the new droplets to shrink and small ones to grow in the formerly-stimulated region. Rather, large droplets are more stable than small ones; over infinite time, Ostwald ripening and droplet coalescence are expected to lead to a single large droplet, properties that are captured in long-timescale simulations of our model (Movie S3; Supplementary Methods). Diffusion, a second process that might act to blur spatial patterns, slows dramatically as droplet size increases, thereby providing a kinetic barrier to erasing spatial memory (Berry et al., 2015). The behavior we observe could have profound implications for a cell, as a transient, locally-applied stimulus could result in a long-term asymmetry in the spatial distribution of protein/RNA droplets, even when individual monomers are able to exchange rapidly in and out of the concentrated phase to interact with other cellular factors (Figure 1G).

### PixELLs: optogenetic control over dissociation of liquid-like protein droplets

Our model suggests that stimulus-dissociated clusters can exhibit long-term spatial memory, but how relevant is this phenomenon at the length- and time-scales of the cell? To address this question, we sought to develop an experimental system to match our modeled scenario: namely, where local stimulation could be used to induce dissociation of protein droplets that assemble spontaneously in the dark. Optogenetic control is ideal for such a study because precise spatial light stimuli can be readily applied and removed. Also, we recently demonstrated that protein phase separation is amenable to optogenetic control by fusing an intrinsically disordered protein region (IDR) to the Cry2 photolyase homology region (PHR) to create OptoDroplets (Shin et al., 2017). In response to light, Cry2 oligomerization nucleates IDR-containing clusters that, over seconds—minutes, grow into micron-scale, liquid-like droplets.

As a starting point for developing an inverse system that confers optogenetic control over droplet dissociation, we turned to two proteins, PixD and PixE, from *Synechocystis* sp. PCC6803 (Masuda et al., 2004; Yuan and Bauer, 2008). PixD and PixE associate in the dark into large multi-subunit complexes (thought to exhibit 10:4 or 10:5 PixD:PixE stoichiometry) that rapidly dissociate into dimers of PixD and monomers of PixE upon blue light stimulation. Upon a shift back to darkness, PixD cycles back to its binding-competent state within seconds to re-form complexes. We reasoned that fusing PixD and PixE to intrinsically disordered protein regions (IDRs) might enable the nucleation of phase-separated droplets in the dark, and that light stimulation might induce the rapid dissociation of these complexes (Figure 2A,B; Figure S2).

**Figure 2.**
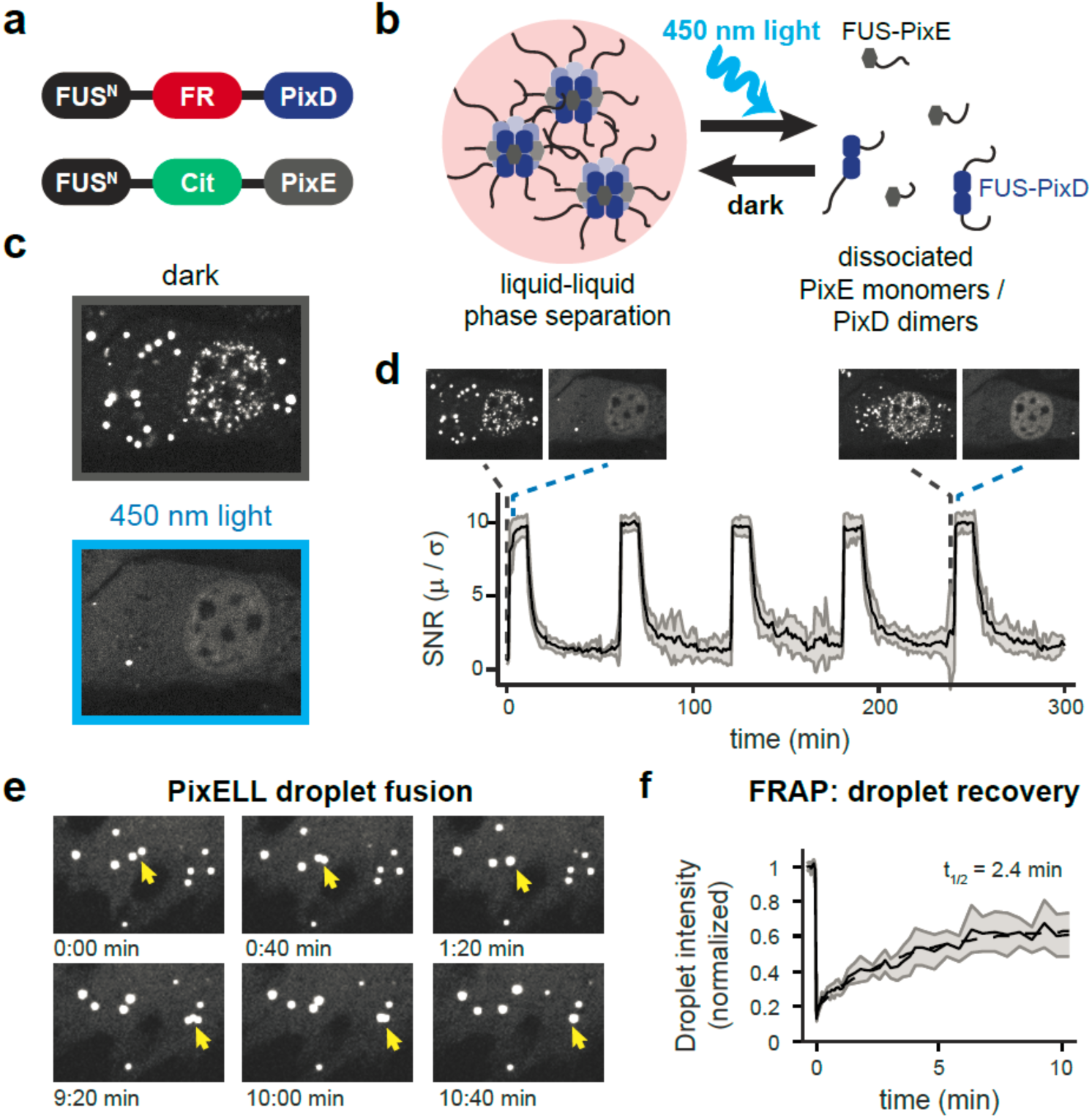
Developing an optogenetic system for spatial control of liquid droplet disassembly. (**a**) Constructs used to create the PixELL optogenetic system and (**b**) schematic of blue light-dissociable intracellular droplets. (**c**) Representative images of intracellular clusters before and after 450 nm light-induced dissociation. (**d**) Signal-to-noise ratio in a cytoplasmic area demonstrate reversible clustering during 5 cycles of dissociation and aggregation. Mean + SEM are shown for 8 representative cells. (**e**) Visualization of two PixELL droplet fusion events. (**f**) Droplet intensity during FRAP experiments indicating photobleaching at t=0 and recovery over 10 min. Mean + SEM are shown for 5 cells, normalized to initial intensity. See Supplemental Methods for details on data collection and quantification.

Indeed, we found that expressing fluorescent FUS^N^-FusionRed-PixD and FUS^N^-Citrine-PixE proteins in NIH3T3 cells led to the formation of micrometer-sized spherical clusters in the dark which dissociated in seconds after blue light stimulation (Figure 2C). Light-controlled clustering was also fully reversible across multiple cycles of photostimulation (Figure 2D; Movie S4). These clusters exhibited hallmarks of phase separation into liquid-like droplets, including droplet fusion, shape relaxation, and fast recovery after photobleaching (Figure 2E,F; Movie S5). Having shown that these IDR-Pix clusters were indeed light-dissociable liquid droplets, we termed them PixELLs (<u>Pix E</u>vaporates from <u>L</u>iquid-like droplets in <u>L</u>ight). In addition to defining the emergent spatiotemporal features of protein phase separation in cells, we expect the PixELL system could serve as a useful optogenetic tool for long-term concentration of proteins into synthetic membraneless organelles (Nakamura et al., 2017; Taslimi et al., 2014) or to sequester and release proteins of interest from subcellular compartments, a phenomenon we demonstrate for light-induced nuclear-to-cytoplasmic transport (Figure S3).

### PixELLs exhibit spatial memory and convert shallow gradients into sharp boundaries

To test if the spatial distribution of PixELLs could encode long-term memory of transient stimuli, we applied and removed a blue light stimulus to subcellular regions of PixELL-expressing cells (Figure 3A; Movie S6). Light exposure induced seconds-timescale dissociation of droplets within the stimulated region, followed by droplet nucleation and growth in unstimulated regions within 10 minutes. Consistent with our model, cells maintained a strongly asymmetric distribution of PixELLs after light stimulus removal, with a sharp boundary between the previously-stimulated and unstimulated cytosolic regions. This persistent asymmetry was evident in both the overall PixELL protein concentration and in the fluorescence intensity of individual droplets (Figure 3B-C). We hypothesized that spatial patterns might be even more striking in subcellular regions where diffusion is slower, such as within the nucleus (Kuhn et al., 2011). Indeed, illuminating the entirety of the cell except for a protected nuclear region enabled us to “draw” precise spatial patterns of droplets in the nucleus that persisted even as the nucleus rotated within the cell (Figure 3D). For both nuclear and cytosolic clusters, asymmetric protein distributions are established within 5-10 min and persist for at least 100 min after the stimulus is removed (Figure S4).

**Figure 3.**
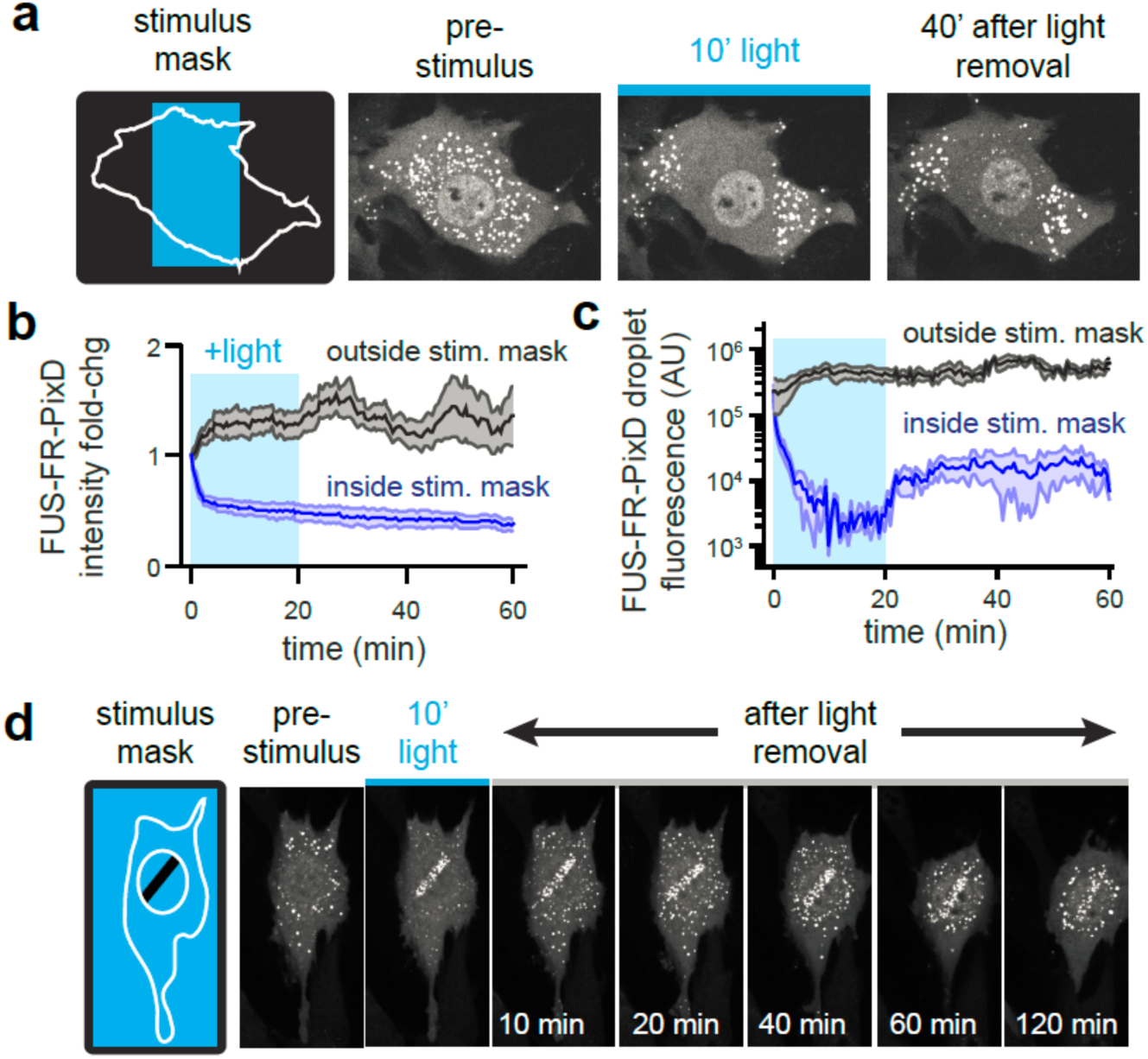
PixELLs exhibit long-term spatial memory of transient stimuli. (**a**) Schematic and images of spatially-restricted 450 nm light stimulation. Fluorescent images of FUS^N^-FusionRed-PixD are shown for cells before, during and after stimulation. (**b**) Cytoplasmic intensity in regions inside and outside the stimulation mask for 4 cells. Mean + SEM are shown. (**c**) Mean cluster size for the cell in **a**, averaged across 5 clusters inside and outside the stimulation area. (**d**) Still images showing long-term memory of a nucleus-localized light stimulus.

We reasoned that PixELLs would also be ideal to test for a second form of spatial information processing: the amplification of a shallow stimulus gradient into a sharp boundary of protein droplets. As is familiar from the of water crossing its freezing point to form ice, phase separation is an inherently all-or-none phenomenon, with the potential to exhibit dramatic physical responses to a small change in an external stimulus (e.g. temperature). Prior theoretical results suggest that this all-or-none effect could also be observed for spatial patterns of intracellular phase separation, where a shallow gradient of a droplet-dissociating stimulus might be converted into a sharp spatial boundary (Lee et al., 2013). Such a scenario is thought to describe P granule dynamics in *C*. *elegans* embryos (Brangwynne et al., 2009). To test this prediction, we applied a linear gradient of 450 nm light intensity to individual PixELL-expressing cells. Indeed, this light gradient induced a sharp boundary of intracellular droplets, converting a shallow stimulus into a switchlike response (Figure 4A-C; Movie S7). This spatial pattern was also retained after stimulus removal, demonstrating both gradient amplification and long-term spatial memory in a single experimental context (Figure 4B). Our results thus demonstrate that protein phase separation is a powerful and versatile way to convert transient, weak biochemical signals into long-lasting spatial patterns in cells.

**Figure 4.**
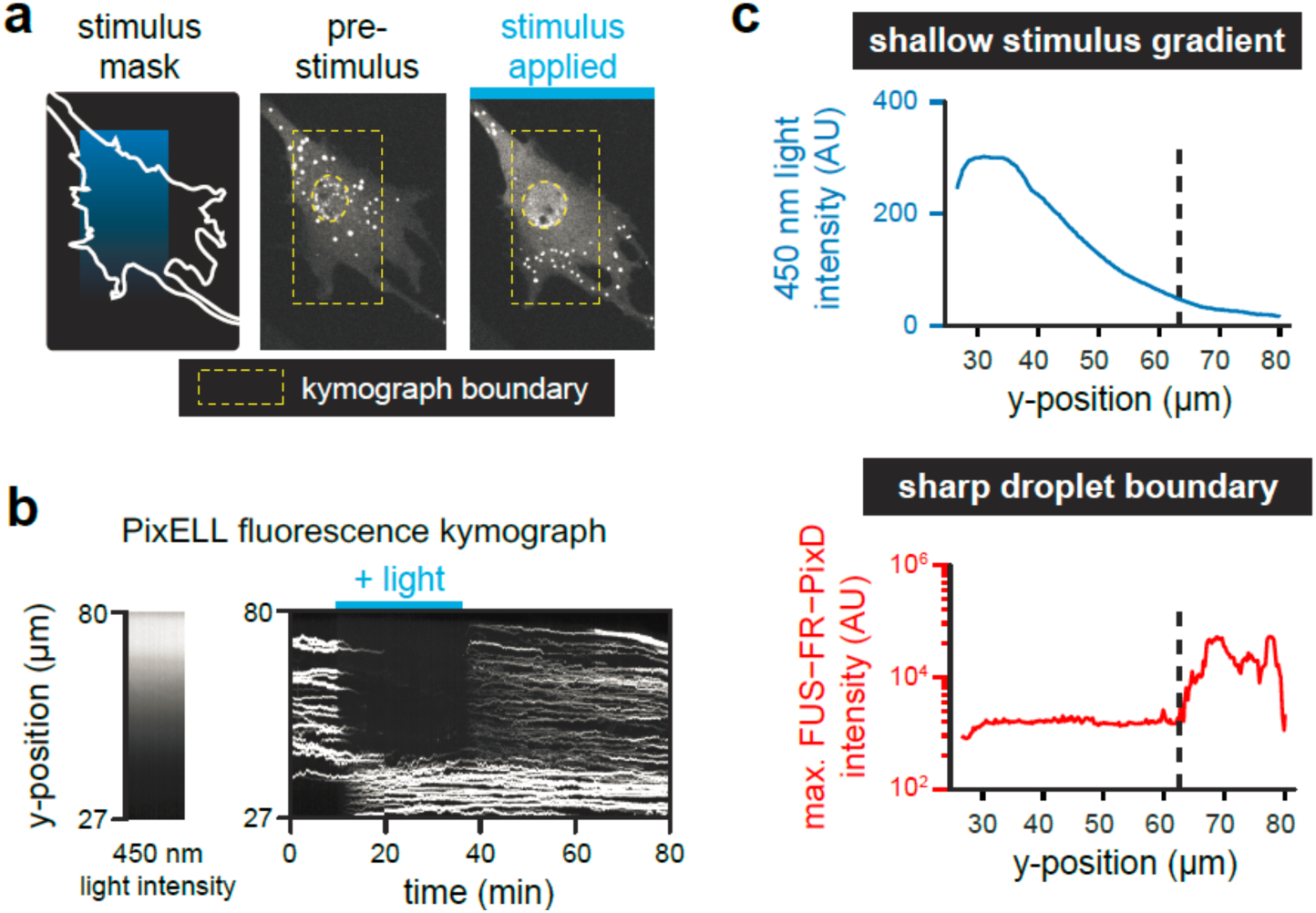
PixELLs amplify shallow stimulus gradients into all-or-none spatial patterns of droplets. (**a**) Gradient stimulation of a PixELL-expressing NIH3T3 cell. Fluorescent images of FUS^N^-FusionRed-PixD are shown for a representative cell stimulated with a linear gradient of light intensity. (**b**) Kymograph of maximum FUS^N^-FusionRed-PixD fluorescence within each row of the yellow box from **a** (right), and median blue light intensity measured within the yellow box from **a** (left). (**c**) Quantification of the kymograph in **b** at 35 min, after spatial light pattern is established. A gradual decrease in 450 nm intensity (top panel; blue curve) elicits a sharp, switch-like transition to form bright FUS^N^-FR-PixD droplets (bottom panel; red curve).

### Membrane-localized optoDroplets also exhibit spatial memory

Our work thus far leaves two important questions unanswered. First, how robust are these phenomena –are they highly dependent on a specific optogenetic tool or cellular context? Second, can the asymmetries in droplet distributions be transmitted to downstream signaling processes to regulate localized cell responses? To address these questions, we set out to probe spatial memory in a distinct and biologically-important context: the membrane-localized signaling clusters formed by activated receptor tyrosine kinases (RTKs). RTKs have been shown to undergo large-scale clustering upon stimulation (van Lengerich et al., 2017) and are often used by cells to drive localized, subcellular responses to external cues (Friedl and Gilmour, 2009). Moreover, optogenetic variants of FGFR1 have been previously shown to drive Erk signaling, cytoskeletal rearrangement and directed migration (Grusch et al., 2014; Kim et al., 2014).

We first sought to adapt our optoDroplet or PixELL system to the plasma membrane to gain control over receptor localization in this subcellular compartment. We found that PixELLs failed to cluster after fusion to an N-terminal myristoylation tag, yet Myr-optoDroplets exhibited robust, light-dependent membrane clustering with a high degree of spatial control (Figure 5A). Local light stimulation of Myr-optoDroplet-expressing cells drove membrane protein clustering only within the illuminated region, and which disassembled within minutes in the dark, consistent with the minutes-timescale half-life of the Cry2 photoactivated state (Figure 5B-C). We noticed that some Myr-optoDroplets exhibited fast, directional motion within the membrane, suggesting active transport along some cytoskeletal components. This directional movement was abolished by treatment with nocodazole but not latrunculin A or a carrier control, suggesting that it is microtubule-dependent (Movie S8). Nevertheless, despite this active transport, the majority of optoDroplets remain localized to the illuminated region over time.

**Figure 5.**
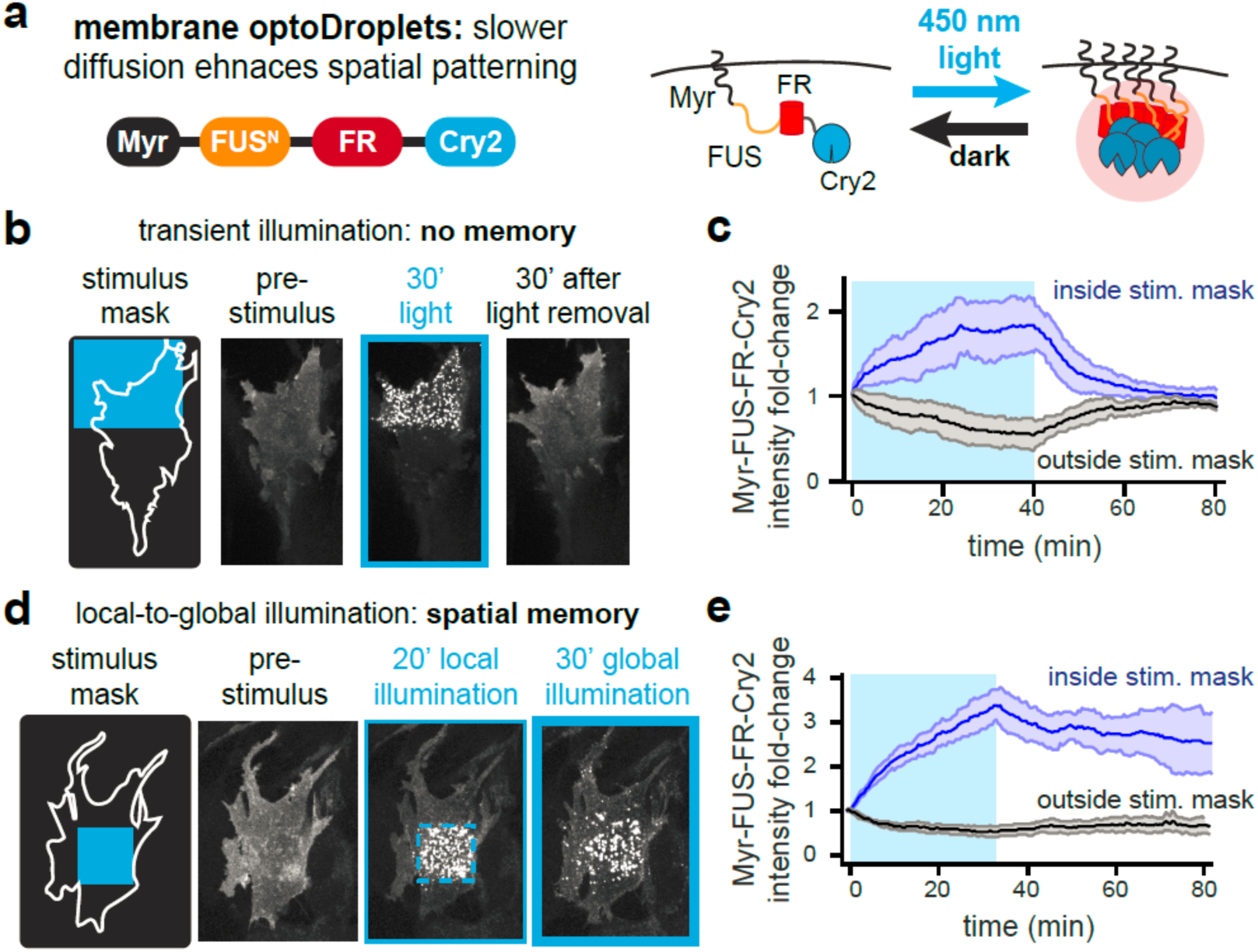
Membrane-localized optoDroplets retain spatial memory of transient stimuli. (**a**) Schematic of Myr-optoDroplet construct and mode of activation. (**b**) Still images of Myr-FUS^N^-FusionRed-Cry2 for a cell exposed to a transient, local 450 nm stimulus. (**c**) Quantification of total intensity for membrane regions inside and outside the stimulus mask, respectively. Mean + SEM are shown for 3 cells. (**d**) Schematic and still images of Myr-FUS^N^-FusionRed-Cry2 localization in the membrane plane for a cell exposed to a local 450 nm stimulus (dashed blue box) followed by global 450 nm illumination. (**e**) Quantification of total intensity in membrane regions inside and outside the stimulus mask, respectively. Mean + SEM are shown for 3 cells.

We next sought to test if membrane clustering might also exhibit the hallmarks of long-term biophysical memory. However, unlike PixELLs, optoDroplet clustering is induced rather than dissociated by light. We have shown that spatial memory requires a cluster-dissociating stimulus, necessitating the use of a new stimulus protocol to test for memory. We reasoned that light-induced clustering could trigger memory formation if an initially local stimulus were then expanded to a global stimulus, a scenario where the *unilluminated* region is thus treated as the localized cluster-dissociating stimulus that is removed upon the shift to global illumination. Such a “local-to-global” stimulus protocol resembles the transition of a migrating cell from a chemoattractant gradient to a high uniform source; in this scenario it may well be advantageous for the cell to preserve memory of the most recent spatial gradient it encountered (Prentice-Mott et al., 2016).

Indeed, we found that this local-to-global stimulus protocol was able to maintain a local pattern of membrane Myr-optoDroplets for at least 1 h after a shift to global illumination (Figure 5D,E; Movie S9). Local illumination induced optoDroplet clustering within 5-10 min, profoundly decreasing the membrane optoDroplet concentration at un-illuminated positions and preventing cluster formation at these positions after the shift to global illumination. Taken together, our data demonstrates that spatial memory is highly robust, operating with similar kinetics in three distinct subcellular compartments (the cytosol, the nucleus and the plasma membrane) and with two optogenetic systems (PixD/PixE-based PixELLs and Cry2-based Myr-optoDroplets).

### FGFR1 droplets can harness spatial memory to drive asymmetric cytoskeletal responses

We finally set out to probe whether the long-term memory encoded in spatial distribution of clusters could be functionally coupled to cell behavior. We first adapted our Myr-optoDroplet construct by fusing it to the cytoplasmic domain of the FGFR1 receptor to create the FGFR1-optoDroplet system (Figure 6A). We observed weaker but qualitatively-similar clustering in FGFR1-optoDroplet cells as compared to Myr-optoDroplets. A limitation on droplet growth could result from FGFR1-mediated phosphorylation of FUS^N^ or Cry2, thereby alter their intrinsic capacity for oligomerization, or from recruitment of additional proteins that decrease optoDroplet association.

**Figure 6.**
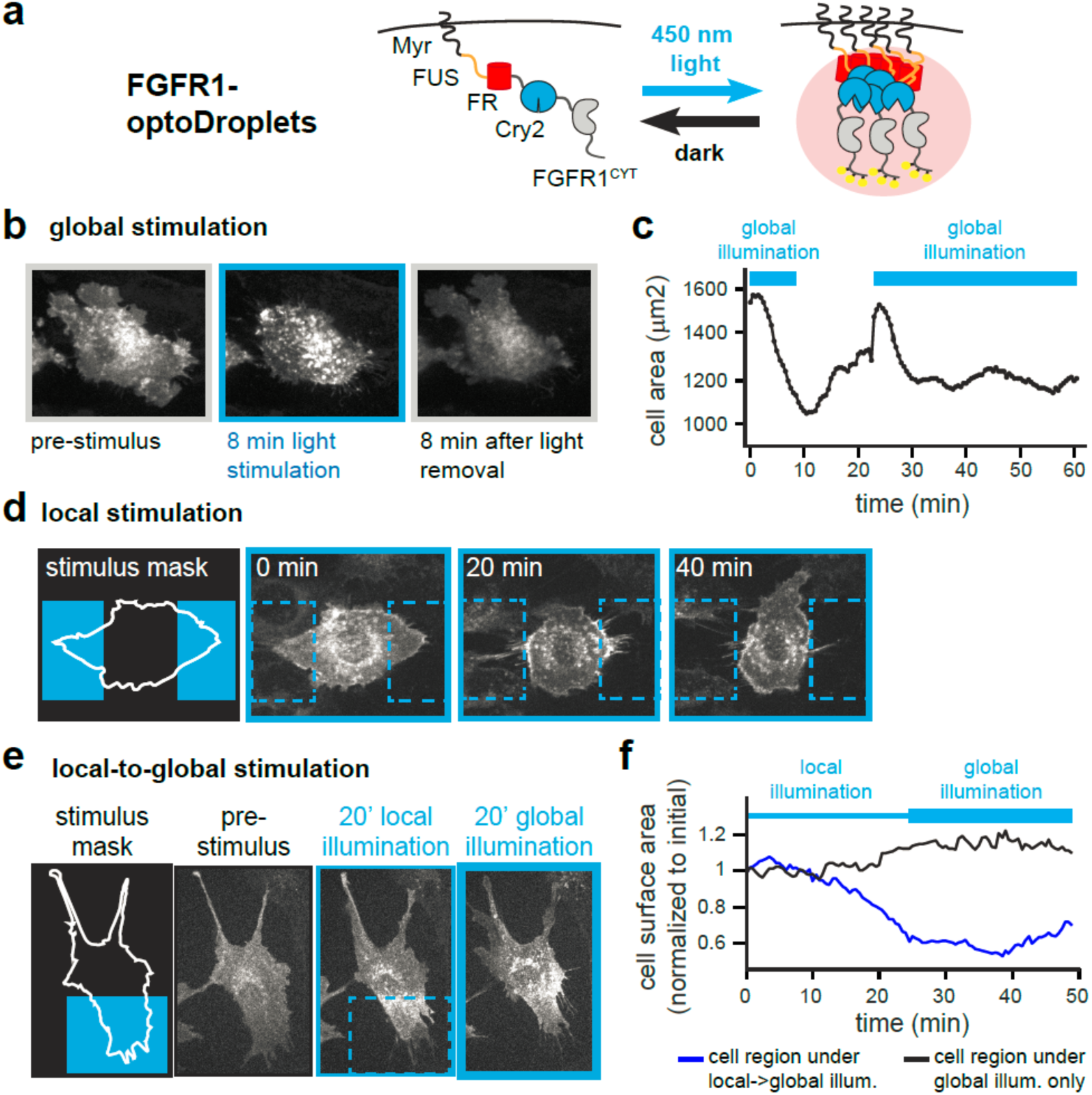
Liquid phase separation drives spatial memory in RTK signaling. (**a**) Schematic showing FGFR1-optoDroplets for inducing RTK clustering and downstream signaling. For all experiments in Fig. 6, Myr-FUS^N^-FusionRed-Cry2-FGFR1 localization is shown. **(b)** FGFR1-optoDroplet cells reversibly “cringe” in response to global blue light stimulation. **(c)** Quantification of change in cell surface area for cell pictured in **b. (d)** FGFR1-optoDroplet cells retract in response to light, ‘avoiding’ a local light stimulus. (**e**) FGFR1-optoDroplet cells exhibit persistent local clustering and cytoskeletal contraction even after a switch to global illumination. (**f**) Quantification of cell surface area within the local-to-global illuminated region (blue box in **e**) and global-only illuminated region (remainder of cell in **e**) during the local-to-global illumination experiment.

We validated that FGFR1-optoDroplets were functional and able to drive potent, light-switchable signaling responses. Illumination led to Erk activation within minutes, as measured by the Erk activity biosensor, ErkKTR, which leaves the nucleus in response to Erk phosphorylation (Figure S5) (Regot et al., 2014). Illuminating FGFR1-optoDroplet cells also elicited a pronounced cytoskeletal response: global “cringing” of entire cells in response to uniform illumination that persisted as long as light was present and was quickly reversed in the dark (Figure 6B-C, Movie S10).

Spatial patterning and long-term memory were also evident in the subcellular distribution of FGFR1-optoDroplets and cytoskeletal activity. Local light stimulation also induced local cell contraction, leading to reorientation of cells to avoid blue light illumination (Figure 6D; Movie S11). In cells subjected to our local-to-global stimulus protocol, FGFR1-optoDroplet cells drove immediate retraction of the plasma membrane within the stimulated region that also persisted upon a subsequent switch to global illumination. Just as observed in the case of Myr-optoDroplets, FGFR1-optoDroplets were concentrated at the initial site of local activation and depleted elsewhere, thereby preventing additional receptor clustering after the shift to global illumination (Figure 6E,F; Movie S12). Notably, the region of the plasma membrane with FGFR1-optoDroplet clusters moved with the retracting protrusion, but a “corset” of contractility was retained on the cell at the same position as these clusters throughout this process. Our results thus show that the spatial memory encoded by protein clusters can be functionally coupled to receptor activation on the plasma membrane, an important spatially-localized cellular response.

## Discussion

Our computational and experimental findings demonstrate that a deceptively simple biophysical system – liquid droplets whose interaction strength is controlled by a spatial stimulus – is sufficient to maintain highly asymmetric, polarized protein distributions in live cells. We also demonstrate that protein phase separation is sufficient to amplify weak spatial stimuli into all-or-none responses, a phenomenon that was previously predicted based on *in vivo* observations of P granule condensation. When considered together, these properties suggest a model where protein condensation plays the role of a highly sensitive ‘memory foam’. Even a weak, transient stimulus – or shallow stimulus gradient – can drive sharp boundaries of protein droplets that persist for an order of magnitude longer than they take to establish. The formation and dissociation of intracellular phase-separated structures thus constitutes a simple and universal mechanism for spatially regulating biological processes.

Although the underlying physics that govern phase separation are well understood, we still know very little about how these processes play out within cells, a scenario that is complicated by the action of complex intracellular processes (e.g. cytoplasmic flow; cytoskeletal assembly/disassembly) as well by potential regulation (e.g. directed transport or regulated disassembly of protein droplets). Here, we show that the phenomenon of spatial memory is remarkably robust, operating within cells on 2-dimensional surfaces (the cell membrane) and 3-dimensional compartments (the cytosol and nucleus). It arises in the context of optogenetic systems from different sources (plant vs. cyanobacteria), and which utilize distinct photosensitive domains (PHR vs BLUF domains).

Our results are reminiscent of the hysteresis observed in classic bi-stable biological systems (Xiong and Ferrell, 2003) but arise through a distinct mechanism. They do not represent a stable steady state formed by the action of positive/negative biochemical feedback loops, as in the case of the spatial patterns that emerge spontaneously from a Turing reaction-diffusion system. Instead, they are rooted in a kinetically-trapped biophysical process: an asymmetric distribution of protein clusters that is unable to mix quickly by diffusion or proceed to the equilibrium of a single connected droplet. Nevertheless, sources of positive and negative feedback are intrinsic to phase separation: as clusters grow, they become more stable and grow still faster (a form of local positive feedback), leading to the depletion of free monomers from solution (a form of long-range negative feedback). It is tempting to speculate that protein phase separation might provide an exceedingly simple and universal way to store spatial information in biological systems, providing a substrate on which to layer more complex biochemical circuits for establishing and maintaining spatial patterns.

## Author contributions

Conceptualization, E.D., A.A.G. and J.E.T.; Methodology, E.D., C.P.B. and J.E.T.; Investigation, E.D., G.U., and J.E.T.; Writing – Original Draft, E.D. and J.E.T.; Writing – Review & Editing, all authors; Funding Acquisition, C.P.B. and J.E.T.; Resources, C.P.B. and J.E.T.; Supervision, J.E.T.

## Acknowledgements

We thank all members of the Toettcher lab for helpful comments. We especially thank Dr. Peter Tonge (Stony Brook University) for sharing DNA constructs and expertise for the PixD and PixE proteins. JET was supported by NIH grant DP2EB024247 and ED was supported by NIH Training Grant T32GM007388. We also thank Dr. Gary Laevsky and the Molecular Biology Microscopy Core, which is a Nikon Center of Excellence, for microscopy support.

## References

Altschuler, S.J., Angenent, S.B., Wang, Y., and Wu, L.F. (2008). On the spontaneous emergence of cell polarity. Nature 454, 886–889.

Banjade, S., and Rosen, M.K. (2014). Phase transitions of multivalent proteins can promote clustering of membrane receptors. eLife 3.

Berry, J., Weber, S.C., Vaidya, N., Haataja, M., and Brangwynne, C.P. (2015). RNA transcription modulates phase transition-driven nuclear body assembly. Proc Natl Acad Sci U S A 112, E5237–5245.

Brangwynne, C.P., Eckmann, C.R., Courson, D.S., Rybarska, A., Hoege, C., Gharakhani, J., Julicher, F., and Hyman, A.A. (2009). Germline P granules are liquid droplets that localize by controlled dissolution/condensation. Science 324, 1729–1732.

Doi, M. (2013). Soft matter physics (Oxford University Press).

Forrest, K.M., and Gavis, E.R. (2003). Live imaging of endogenous RNA reveals a diffusion and entrapment mechanism for nanos mRNA localization in Drosophila. Current biology: CB 13, 1159–1168.

Freeman Rosenzweig, E.S., Xu, B., Kuhn Cuellar, L., Martinez-Sanchez, A., Schaffer, M., Strauss, M., Cartwright, H.N., Ronceray, P., Plitzko, J.M., Forster, F., et al. (2017). The Eukaryotic CO2-Concentrating Organelle Is Liquid-like and Exhibits Dynamic Reorganization. Cell 171, 148–162 e119.

Friedl, P., and Gilmour, D. (2009). Collective cell migration in morphogenesis, regeneration and cancer. Nature reviews Molecular cell biology 10, 445–457.

Gierer, A., and Meinhardt, H. (1972). A theory of biological pattern formation. Kybernetik 12, 30–39.

Goldstein, B., and Macara, I.G. (2007). The PAR proteins: fundamental players in animal cell polarization. Developmental cell 13, 609–622.

Grusch, M., Schelch, K., Riedler, R., Reichhart, E., Differ, C., Berger, W., Ingles-Prieto, A., and Janovjak, H. (2014). Spatio-temporally precise activation of engineered receptor tyrosine kinases by light. The EMBO journal 33, 1713–1726.

Kim, N., Kim, J.M., Lee, M., Kim, C.Y., Chang, K.Y., and Heo, W.D. (2014). Spatiotemporal control of fibroblast growth factor receptor signals by blue light. Chemistry & biology 21, 903–912.

Kloc, M., and Etkin, L.D. (2005). RNA localization mechanisms in oocytes. J Cell Sci 118, 269–282.

Kuhn, T., Ihalainen, T.O., Hyvaluoma, J., Dross, N., Willman, S.F., Langowski, J., Vihinen-Ranta, M., and Timonen, J. (2011). Protein diffusion in mammalian cell cytoplasm. PloS one 6, e22962.

Landau, D.P., and Binder, K. (2014). A guide to Monte Carlo simulations in statistical physics (Cambridge university press).

Lee, C.F., Brangwynne, C.P., Gharakhani, J., Hyman, A.A., and Julicher, F. (2013). Spatial organization of the cell cytoplasm by position-dependent phase separation. Physical review letters 111, 088101.

Masuda, S., Hasegawa, K., Ishii, A., and Ono, T.A. (2004). Light-induced structural changes in a putative blue-light receptor with a novel FAD binding fold sensor of blue-light using FAD (BLUF); Slr1694 of synechocystis sp. PCC6803. Biochemistry 43, 5304–5313.

Nakamura, H., Lee, A.A., Afshar, A.S., Watanabe, S., Rho, E., Razavi, S., Suarez, A., Lin, Y.C., Tanigawa, M., Huang, B., et al. (2017). Intracellular production of hydrogels and synthetic RNA granules by multivalent molecular interactions. Nat Mater.

Prentice-Mott, H.V., Meroz, Y., Carlson, A., Levine, M.A., Davidson, M.W., Irimia, D., Charras, G.T., Mahadevan, L., and Shah, J.V. (2016). Directional memory arises from long-lived cytoskeletal asymmetries in polarized chemotactic cells. Proceedings of the National Academy of Sciences of the United States of America 113, 1267–1272.

Regot, S., Hughey, J.J., Bajar, B.T., Carrasco, S., and Covert, M.W. (2014). High-sensitivity measurements of multiple kinase activities in live single cells. Cell 157, 1724–1734.

Sailer, A., Anneken, A., Li, Y., Lee, S., and Munro, E. (2015). Dynamic Opposition of Clustered Proteins Stabilizes Cortical Polarity in the C. elegans Zygote. Developmental cell 35, 131–142.

Shin, Y., Berry, J., Pannucci, N., Haataja, M.P., Toettcher, J.E., and Brangwynne, C.P. (2017). Spatiotemporal Control of Intracellular Phase Transitions Using Light-Activated optoDroplets. Cell 168, 159–171 e114.

Shin, Y., and Brangwynne, C.P. (2017). Liquid phase condensation in cell physiology and disease. Science 357.

Su, X., Ditlev, J.A., Hui, E., Xing, W., Banjade, S., Okrut, J., King, D.S., Taunton, J., Rosen, M.K., and Vale, R.D. (2016). Phase separation of signaling molecules promotes T cell receptor signal transduction. Science 352, 595–599.

Taslimi, A., Vrana, J.D., Chen, D., Borinskaya, S., Mayer, B.J., Kennedy, M.J., and Tucker, C.L. (2014). An optimized optogenetic clustering tool for probing protein interaction and function. Nat Commun 5, 4925.

Turing, A.M. (1990). The chemical basis of morphogenesis. 1953. Bulletin of mathematical biology 52, 153-197; discussion 119-152.

van Lengerich, B., Agnew, C., Puchner, E.M., Huang, B., and Jura, N. (2017). EGF and NRG induce phosphorylation of HER3/ERBB3 by EGFR using distinct oligomeric mechanisms. Proceedings of the National Academy of Sciences of the United States of America 114, E2836–E2845.

Xiong, W., and Ferrell, J.E., Jr. (2003). A positive-feedback-based bistable ‘memory module’ that governs a cell fate decision. Nature 426, 460–465.

Yuan, H., and Bauer, C.E. (2008). PixE promotes dark oligomerization of the BLUF photoreceptor PixD. Proceedings of the National Academy of Sciences of the United States of America 105, 11715–11719.

