## Supplementary Materials for "Protein phase separation provides long-term memory of transient spatial stimuli"

**This PDF file includes:**

Supplementary Methods
Figs. S1 to S5
Captions for Movies S1 to S12

### Supplementary Methods

#### Computational model

##### *Description of the model*

Our computational model was intended to capture the basics of diffusion, association and dissociation of self-associating "proteins" in a simple and minimal context, where proteins were modeled as single elements on a 2 dimensional grid. In our model, proteins move randomly across the grid. Upon occupying grid spaces with neighboring spaces that are also occupied, proteins experience identical binding interactions with each neighbor that decreases the probability of movement away from that position. Interactions are assumed to be maximally conservative: if a protein moves from one grid space to an adjoining one where some neighbors are shared, it is assumed that these binding interactions are not broken so they do not contribute to the penalty. For all simulations, we used a 50 x 100 grid with reflective boundary conditions that was populated by a random initial distribution of 700 proteins monomers.

We define a contiguous set of interacting proteins as a "cluster". Such clusters may exhibit liquid-like or solid-like properties depending on the interaction strength and other system parameters.

Our system thus admits three kinds of processes with the following rates (also see Figure S1A):

- **Diffusion**, modeled as movement of a protein from its position on the lattice to a neighboring, unoccupied position. Without loss of generality, we take the rate of diffusive movement as the reference timescale for our simulations, setting the rate constant $d = 1$ in all cases.
- **Exchange** of proteins between two neighboring grid positions. This exchange reaction can also be thought of as diffusion within the clustered phase. We assume that exchange within the clustered phase is half as likely as diffusion to an empty position (i.e. $e = 0.5$). Because exchange involves movement of *two* proteins, halving this rate leads to a comparable timescale of overall protein movement within both phases.
- **Unbinding**, modeled as the breaking of neighboring interactions upon movement away from a grid location. We assume that each protein-protein interaction contributes an interaction energy $\Delta E$, so an unbinding reaction proceeds according to the rate constant $k = k_0 \exp\left(-\frac{\Delta E \cdot n_{lost}}{\theta(x,y,t)}\right)$, where $n_{lost}$ is the number of neighbors whose contacts are broken upon moving away from the prior grid position and $\theta(x,y,t)$ is a stimulus parameter that we can vary at each time point and grid position. The stimulus $\theta$ can be thought of as a temperature-like scaling factor on the interaction energy. For high values of $\theta$, interactions are relatively weak and the system remains in a diffuse state. As $\theta$ is lowered, interaction energies become stronger and the system enters different forms of an aggregated state. The parameter $k_0$ is a constant that is related to the off-rate for breaking a single interaction. For all simulations, we took $\Delta E = 1$ (without loss of generality because the scale of $\theta(x,y,t)$ is set by our input) and $k_0 = 1$ so that unbinding in the limit where no neighbors are lost is identical to diffusion.

*Simulating the model*

We simulated random trajectories for this 2D diffusion/aggregation system using a rejection kinetic Monte Carlo approach (rKMC). rKMC is quite closely related to the Gillespie algorithm that is often used for stochastic chemical systems, and which provably achieves the same results (Serebrinsky, 2011). Rejection kinetic Monte Carlo is highly efficient and straightforward to implement for systems with many reactions under conditions where there is a well-defined "fastest" reaction rate (in our case, that of diffusion), and where a large number of the possible reactions in each configuration proceed at or near this fastest rate (in our case, diffusion and exchange are highly likely be picked, as only the particles at the interface between two phases can undergo unbinding).

In brief, the algorithm proceeds as follows. We pick a random protein on the lattice and random direction for it to move. Such a movement in any direction corresponds to one of the three reactions as defined above (diffusion, exchange or unbinding). If a reaction rate is equal to 1 (which is our fastest rate, and that of diffusion to an empty square), it is automatically accepted. Otherwise, a random number $r_3$ is rolled on the interval [0,1) and the reaction is accepted in proportion to its rate (e.g. if its rate is greater than $r_3$). After each iteration the time *t* is incremented so that $t_i = t_{i-1} + \tau$, where $\tau = 1/N_{rxns} \log(1/r_4)$. In this formula $N_{rxns}$ is the total number of possible reactions in the system and $r_4$ is a fourth random number on the interval [0,1).

Note that $N_{rxns}$ can easily be defined because all possible reactions are uniquely identified with movement from an occupied lattice position to any adjoining lattice position. Thus, the total number of reactions is equal to the sum of all neighboring lattice positions for each occupied lattice position (for the reflective boundary conditions we implement, all middle lattice positions have 8 neighbors; all edge positions have 5; each corner has 3). (Note: MATLAB code implementing our approach is provided as Supplementary Code.)

*Analyzing spatial properties from simulations*

To compute parameters of individual clusters (e.g. their size or solidity), we first identified all connected components at each simulation timepoint using the standard MATLAB image processing function `bwconncomp`. From this list of all connected components in the image we then computed various properties, including the area and solidity of each connected component. Area is useful as a straightforward measure of cluster size. Solidity is defined as the area of a cluster divided by the convex area in which it can be enclosed; it provides a measure of whether clusters are filled, as expected if they are able to relax to a shape that minimizes surface area. Additionally, for each spatial stimulus simulation we measured the total number of proteins in the stimulated and unstimulated regions.

*Performing computational FRAP experiments:*

FRAP analysis was performed in the model by saving the X-Y position of each monomer, and by tracking a single chosen "cluster" over time from an initial time $t_0$ after clusters were established. Fluorescence "recovery" was then associated with the number of monomers within the cluster that had been exchanged for monomers not initially present.

We first identified all the monomers in a chosen cluster by selecting a particular connected component for analysis. For every subsequent timepoint we tracked the cluster's position on the 2D lattice using morphological reconstruction (implemented by the MATLAB function `imreconstruct`), where the cluster at the prior timepoint was taken as a marker image.

We computed two quantities at each timepoint: $n_i$, the number of monomers in the cluster at the i$^{th}$ timepoint, and $o_i$, the intersection between the initial set of monomers in the cluster and the set of monomers at the i$^{th}$ timepoint. We then measured the percent recovery using the formula:

$$f_i = \left(\frac{n_i - o_i}{n_i}\right)\left(\frac{N}{N - n_0}\right)$$

In this formula, the degree of recovery is captured in the first term; the second term is a normalization factor that accounts for the fraction of the total monomer pool that is "bleached" and would be expected to re-enter the cluster.

Model results

*The model capture phase separation as a function of interaction strength*

We first simulated our model for different values of the interaction temperature $\theta$ applied as a uniform global input, i.e. at all lattice positions and times (Figure S1b). We simulated $10^5$ reactions at each temperature (which corresponded to a total simulation time of $T \approx 18$ normalized time units at all temperatures). We found that over a narrow range of $\theta$ values, centered approximately at $\theta = 1$, the system became organized into a phase-separation-like state. At this state we observed a considerable fraction of subunits organized into aggregates, as well as subunits that persisted in the monomeric state. This state was characterized by fast recovery after photobleaching and substantial shape relaxation of the aggregates, both hallmarks of liquid-like behavior (Figure S1C,D). We thus used $\theta = 1$ to simulate the liquid-like state in all subsequent spatial stimulus experiments.

At progressively lower values of $\theta$ the cluster size continued to increase until virtually all subunits were contained within the aggregated phase. After further decreases in $\theta$ we observed a transition to a state with arrested dynamics: cluster size *decreased* from its maximum size because dissociation from a cluster became highly unlikely, leading to a large number of small, spatially-separated clusters. In our simulation this state persisted for long times, although it is likely that a more complex simulation accounting for diffusive or convective transport of entire clusters would serve to induce further aggregation by coalescence.

*A localized decrease in interaction strength leads to local, memoryless aggregation*

We next used the model to probe how a localized stimulus would affect aggregation/ dissolution (Figure S1E; Figure 1B). To model a stimulus-induced local increase in aggregation, we initially evolved the system with $\theta = 2$ at all positions, then transiently dropped the "temperature" to $\theta = 1$ on the right-half of the grid (i.e. for columns 51-100), leaving $\theta = 2$ on the left-half of the grid (columns 1-50) (Figure S1f). Each stimulation regime – initial equilibration, transient stimulation, and stimulus removal – was performed for $2x10^6$ simulation steps, corresponding to approximately 360 normalized time units (this total time varies slightly between simulations based on the stochastic nature of reactions in the kinetic Monte Carlo framework). We found that this stimulus protocol led to transient cluster formation and droplet growth, followed by quick reversal upon stimulus removal (Figure S1F-H). We also tested the converse experiment, meant to represent a stimulus-induced local *decrease* in association strength (Figure 1B). In this case, the system was evolved at the stimulus strength $\theta = 1$ everywhere until it was transiently increased to $\theta = 2$ on the right-half of the grid. This scenario, described in the main text, led to a persistent asymmetry of cluster formation (Figure 1B-E).

*Long-term model simulations show Ostwald ripening and validate the stability of a single, connected droplet*

For phase separating systems, it is expected that the thermodynamic steady state is the formation of a single, connected droplet, a familiar like the separation of oil and vinegar in salad dressing. Approach to this state can be driven by multiple processes, including collisions and fusion between distinct droplets and Ostwald ripening, where large droplets grow at the expense of smaller ones by exchanging monomers that diffuse between droplets. However, the rate of approach to this state can be slow, leading to the quasistable appearance of long-lived, large droplets.

We set out to test whether our model exhibits classic phase separation behaviors: Ostwald ripening, kinetically-trapped states with multiple large droplets, and an equilibrium state defined by a single large droplet. To do so we ran long simulations of at least $10^7$ individual reactions, corresponding to ~2,000-3,000 time units (time units are comparable between all simulations). The results, presented in Movie S3, show that indeed Ostwald ripening can occur, where an initial distribution of ~15 droplets slowly ripens into 5 larger droplets over time, but these 5 droplets remain stable for at least 1,000 time units. In addition, we found that when the model is initialized with a single cluster, its shape relaxes to become approximately circular but no new droplets are formed over the entire course of the simulation. To perform this simulation we initialized the 50 x 100 grid with 700 molecules located in a single rectangle in the center of the grid.

These long-timescale observations compare favorably to those we find in cells using the PixELL and OptoDroplet systems, where multiple large droplets persist over long periods of time. Indeed, we never observe coalescence into a single intracellular droplet, suggesting that the final fusion events or Ostwald ripening occur extremely slowly, or that other active processes (e.g. the regulated disassembly of large droplets) prevent this equilibrium state from being attained.

<u>Plasmids:</u>

All plasmids were constructed using inFusion cloning (Clontech) to ligate in a PCR product to a pHR vector that was opened using either backbone PCR or restriction digest.

*PixELL Constructs*:

To create pHR-FUS$^{N}$-FusionRed-PixD, pHR-FUS$^{N}$-Citrine-PixE and pHR- FUS$^{N}$ -irFP-PixE (see Figure S2) we used backbone PCR off of the plasmids from Shin et al. 2017 to replace the mCherry Fluorescent protein (FP) with the respective FPs. We then conducted backbone PCR on the FUS$^{N}$ -FP-Cry2 to replace the Cry2 with PCR products encoding PixD or PixE. DNA encoding PixD came from plasmids used in Gil et al. 2017, while plasmids encoding PixE DNA were gifts of the Tonge lab.

*Versatility of PixELL System:*

We wished to test the versatility of the PixELL system to check its usefulness for future applications, as certain optogenetic protein support fusions in specific orientations. For PixELLs to be most useful the Pix proteins should support both N-terminal fusions, as shown in the main text, as well as C-terminal fusions. Thus, we tested whether C-terminal fusions of the Pix proteins to FUS$^{N}$-FP could support clusters (Figure S2A-B). To create pHR-PixD-FUS$^{N}$-FusionRed and pHR-PixE-FUS$^{N}$-irFP we digested an empty pHR plasmid with BamHI and NotI and inserted FUS$^{N}$-FP into that vector. We then cut the pHR- FUS$^{N}$-FP plasmid with BamHI to insert PixD or PixE. We did not use pHR-PixE-FUS$^{N}$-irFP or pHR- FUS$^{N}$-irFP-PixE for main text figures, as for unknown reasons these plasmids led to the formation of many more non-light sensitive clusters than the citrine constructs (Figure S2).

*Minimum requirements for PixELL System:*

To see the minimum sequence necessary for cluster formation we removed FUS$^{N}$ from either PixD or PixE (Figure S2C-D). To create these plasmids, pHR-irFP-PixE and pHR-FusionRed-PixD constructs, we used backbone PCR to remove FUS$^{N}$ and the FP and used PCR and inFusion to replace the FP. Finally, we tested whether FUS$^{N}$-FP-PixD or FUS$^{N}$-Citrine-PixE alone could support cluster formation, and they were unable to (Figure S2 E,F). Therefore, the full construct used in the main text is the minimum necessary to generate clusters, limiting the size of additional components one could add to the PixELL plasmids.

*Using PixELLs to Sequester Proteins of Interest:*

We wished to test whether PixELL could be used to sequester proteins of interest in different cellular compartments. We reasoned that by localizing one Pix component to the nucleus, the other Pix component would be sequestered to the nucleus in the dark due to its obligate interaction and formation of clusters. However, in the light we expect the clusters to dissociate, releasing the sequestered Pix protein and allowing it to translocate to the cytosol. Given the reversibility of the PixELL system (Figure 2D), we also reasoned that this sequestration and release would be reversible, making PixELL an incredibly useful tool for reversible sequestration of proteins of interest.

To test this hypothesis, we made pHR-NLS- FUS$^{N}$-Cit-PixE by conducting a BamHI digest to open the pHR-Fus-Cit-PixE plasmid and then PCR amplified a known NLS (MAPKKKRKVRYPAFLYKVA) and used inFusion to ligate the two together. We expressed this plasmid with pHR- $FUS^N$-FusionRed-PixD in NIH-3T3 cells to conduct the translocation experiments in Figure S3.

*Making Membrane-Tethered OptoDroplets:*

We previously showed that local stimulation of cytosolic optoDroplets leads to diffusion of photoactivated monomers and droplet nucleation over a broad cellular area (Shin et al., 2017). We reasoned that in contrast, the reduced diffusion of membrane-optoDroplets should led to sharp patterns of droplets in a local illuminated region. To make constructs that could check this prediction we made myristoylated (MYR) optoDrop. To make pHR-MYR-$FUS^N$ -FusionRed-Cry2 we conducted an MluI digest on the pHR-Fus-FusionRed-Cry2 to remove the Fus-FusionRed sequence. We then added the standard myrostylation sequence (MGSSKSKPKDASQRRR) to PCR primers for an insert that amplified the $FUS^N$ and FusionRed sequence and conducted an InFusion ligation with the MluI digested pHR- $FUS^N$-FusionRed-Cry2.

*Plasmids for FGFR1 experiments:*

To create pHR-Myr-$FUS^N$-FusionRed-Cry2– FGFR1 we used human FGFR1 DNA that we obtained from the R777-E083 Hs.FGFR1 plasmid, which was a gift from Dominic Esposito (Addgene plasmid # 70367). We PCRed the cytoplasmic domain of FGFR1 (amino acids 398-822) as had been done previously (Kim et al., 2014) and ligated it to NotI digested pHR-Myr- $FUS^N$-FusionRed-Cry2 plasmids.

DNA encoding the Erk-Kinase Translocation Reporter (KTR) was a gift from Markus Covert (Regot et al., 2014). We digested pHR-IRFP with BamHI and ligated the ErkKTR PCR product to create pHR-ErkKTR-IRFP.

<u>Cell culture</u>

NIH 3T3 mouse embryonic fibroblasts were grown in DMEM supplemented with 10% FBS, 1% L-Glutamine, and Pen/Strep. Cells were maintained on Thermo Scientific Nunc Cell Culture Treated Flasks with Filter Caps and grown at 37 C with 5% $CO_2$.

<u>Lentivirus production and transduction</u>

Lentivirus was produced as per the protocol we described previously (Toettcher et al., 2013). Briefly, Lenti-X 293T cells were plated in a 6-well plate at 40% confluency and co-transfected with the appropriate pHR expression plasmid and lentiviral packaging plasmids (pMD2.G and p8.91 – gifts from the Trono lab) using Fugene HD transfection reagent. Viral supernatants were collected 2 days after transfection and passed through a 0.45 mm filter.

NIH 3T3 cells to be infected with lentivirus were plated in a 6 well dish at 20%–40% confluency. After adherence to the plate, 500 μl of filtered virus were added to the cells as was 50 μl of 1 M HEPES. 24 h post-infection, viral media was replaced with normal growth media and cells were imaged at least 48 h after infection to allow time for integration and expression.

Cell preparation for imaging

For all imaging experiments cells were plated on black-walled, 0.17 mm glass-bottomed 96 well plates (In Vitro Scientific). Prior to cell plating, glass was pretreated with a solution of 10 µg/mL fibronectin in phosphate buffer saline (PBS) for 20 min. NIH-3T3 cells were allowed to adhere for at least 2 hours in our supplemented DMEM. Just prior to imaging 50 µL of mineral oil was added to the top of each well to stop evaporation (Toettcher et al., 2011).

Time-lapse microscopy

Cells were maintained at 37C with 5% CO2 for the duration of all imaging experiments. Confocal microscopy was performed on a Nikon Eclipse Ti microscope with a Prior linear motorized stage, a Yokogawa CSU-X1 spinning disk, an Agilent laser line module containing 405, 488, 561 and 650 nm lasers, an iXon DU897 EMCCD camera, and a 60X oil immersion objective lens.

Optogenetic stimulation hardware

For microscopy experiments, cells were imaged with the 561 nm laser to image $FUS^N$-FusionRed-PixD in PixELL cell lines, and Myr-$FUS^N$-FusionRed-Cry2 in the optoDroplet cell lines. A 450 nm LED light source (XCite XLED1) was used for all spatial blue light stimulation experiments of both the PixELL and optoDroplet systems, and either the same 450 nm LED light source or 488 nm laser illumination was used for all global experiments. Light from the XLED1 system was delivered through a Polygon400 digital micromirror device (DMD; Mightex Systems) to control the temporal dynamics of light inputs. We applied specific spatial patterns to an image by drawing ROIs within the Nikon Elements software package. To attenuate 450 nm light to appropriate levels, we dithered the DMD mirrors to apply light 10% of the time, and set our 450 nm LED to 5% of its maximum intensity.

For the gradient stimulation experiment in Figure 4, we used a "gradient ROI" in NIS-Elements that allowed a gradient of 0-10% of the light to pass through from left to right. The still images of Figure 4 were rotated 180 degrees from Movie S7 to make the data easier to represent as a kymograph.

For the Myr- and FGFR1-optoDroplet experiments in Figure 5 and Figure 6, where light moved from a local to global pattern, we began the experiment using an ROI to illuminate the region of interest. After the spatial pattern of activation was established, we quickly paused the experiment to resize the ROI so that it would cover the whole field of view, thus switching to global stimulation for the remainder of the experiment.

Finally, for the long term stimulation experiments shown in Figure S4, we stimulated multiple cell positions in the same acquisition, unlike in the earlier experiments, where single positions received constant blue light. Therefore, we had to increase the DMD and LED intensity so that cells received more intense light for a shorter period of time. To do so, we set the DMD mirrors set to a 50% duty cycle and LED power at 50% of maximum intensity.

FGFR1 imaging:

To confirm our FGFR1-optoDroplet construct was functional we measured Erk activity using the ErkKTR (Regot et al., 2014), as Erk activity was shown to respond to a previous optogenetic FGFR1 construct (Kim et al., 2014). We measured KTR presence in the nucleus and cytoplasm in multiple positions for FGFR1-optoDroplets, by imaging with the CSU561 (to confirm that construct would cluster in blue light) and CSU650 (to image ErkKTR localization) and stimulated with DMD light at 40% LED power and with a 60% ROI. Positions were stimulated and imaged once every 30 seconds (Figure S5).

FRAP experiments

FRAP experiments were performed in the Nikon Microscopy Core imaging facility at Princeton University on a point-scanning confocal (A1R-Si on a Nikon Ti-E microscope chassis). Bleaching was performed by applying 7.5% of the maximum power from our 561nm laser on a single cluster. We found that this light was powerful enough to photobleach FusionRed fluorescence but not sufficiently intense to induce optogenetic stimulation and PixELL dissociation. Images were captured pre- and post-bleach using confocal imaging with the 561 nm laser at 0.4% power.

Drug Additions:

For cytoskeletal perturbation experiments, all drugs were reconstituted to 1 mg/ml concentrations in DMSO. For the experiments shown in Movie S8, DMSO was diluted to 0.5% in full media (representing the maximum final DMSO concentration used in any drug treatment), while Latrunculin A was diluted to 5µg/ml in full media and Nocodazole to 2.5µg/ml. 20µl of each solution were added to NIH-3T3 cells in a 100µl of full media in 96 well glass bottom plate 16hrs before imaging.

Image analysis

*Obtaining properties of cellular regions*

All image analysis was performed in ImageJ. First, appropriate nuclear or cytoplasmic regions were tracked over time by hand annotation. We then measured properties of each annotated region at each timepoint, including the mean and standard deviation of pixel intensities, the area of the region, and its integrated intensity (e.g. area * mean). For measuring overall protein redistribution we used the integrated intensity over large, equally-sized cytoplasmic areas inside and outside the stimulation region.

For measuring kinetics of droplet assembly/disassembly, we found that the signal to noise ratio (the inverse of the coefficient of variation) to be an excellent metric that spanned a reproducible range even for cells with different PixELL expression levels. The signal-to-noise ratio is defined as $SNR = \mu / \sigma$ , where $\mu$ is the pixel-by-pixel mean intensity within the region and $\sigma$ is the standard deviation. In particular, we found that the coefficient of variation $CV = \sigma / \mu$ took on large values that could vary substantially between cells with a high degree of droplet formation (or even between different regions of the same cell); by inverting these large numbers, the signal-to-noise ratio compressed their differences and led to reproducible measurements of the kinetics of droplet formation. We thus used the SNR to describe the kinetics of droplet formation in Figure 2.

For some analyses (e.g. FRAP photobleaching recovery; cluster size over time) we analyzed the intensity of individual droplets over the course of a timelapse acquisition. In these cases, we annotated an individual cluster by hand using the ImageJ 'measure' tool. From there we developed a Matlab script that (a) identifies the XY location of the peak intensity in the annotated region at each frame of the time series, (b) fits a 2-dimensional Gaussian to the region $a \cdot \exp\left( -\frac{(x-x_0)^2 + (y-y_0)^2}{2c^2} \right) + b$ , and (c) calculates the integrated area under the fit Gaussian as the burst intensity $I = 2\pi ac^2$.

Figure 6 shows measurements of the sizes of FGFR1-optoDroplet cells undergoing light-induced contraction. To perform these analyses we took advantage of the fluorescence of the optoDroplet construct, which permitted us to segment cells from background by simply applying a threshold. Noisy bright pixels were excluded by a binary opening operation, any dark pixels in the interior of the image were filled in by a morphological hole-filling operation, and finally any remaining noisy regions were excluded by keeping only the biggest connected component at each time point. We then measured the total cell footprint area by measuring the number of pixels in the binary mask at each time point, and converting to units of $\mu m^2$ using the pixel-to-distance calibration of our microscope. For some analyses we further subdivided and tracked the cell footprint area in both the illuminated and non-illuminated regions.

#### Supplementary Figure 1

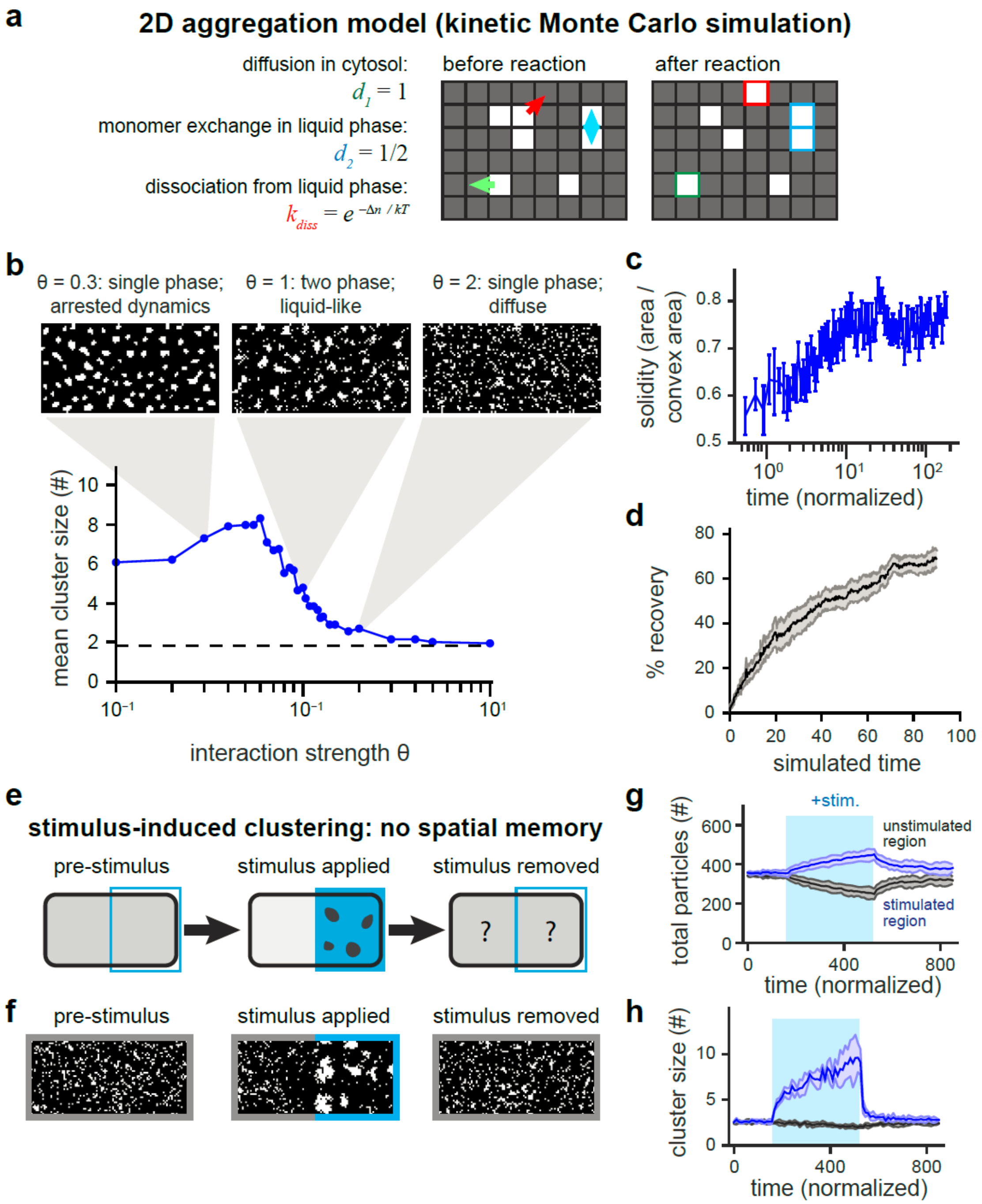


**Figure S1, related to Figure 1. Interrogating a simple computational model of phase separation.** (**a**) Schematic of different reactions in our Monte Carlo simulation. (**b**) Scan of mean cluster size as a function of the protein-protein association strength θ. Inset images show 2D grid after $10^5$ iterations of the kinetic Monte Carlo (~18 time units). (**c**) Solidity of clusters (mean + SD for 10 clusters) shown during droplet formation for the $\theta = 1$ value of interaction strength. A solidity of 1 would indicate that each aggregate fills its convex hull. (**d**) Simulated FRAP recovery for clusters formed in response to the $\theta = 1$ value of interaction strength. (**e**) Schematic of stimulus application and removal. (**f**) Representative images of 2-D lattice showing local clustering during stimulation, with diffuse subunits before and after stimulation. (**g,h**) Total number of subunits (**g**) and mean cluster size (**h**) in the stimulated (right half of lattice) and unstimulated (left half of lattice) regions before, during and after local stimulation. In every panel, error bars show mean + SD.

#### Supplementary Figure 2

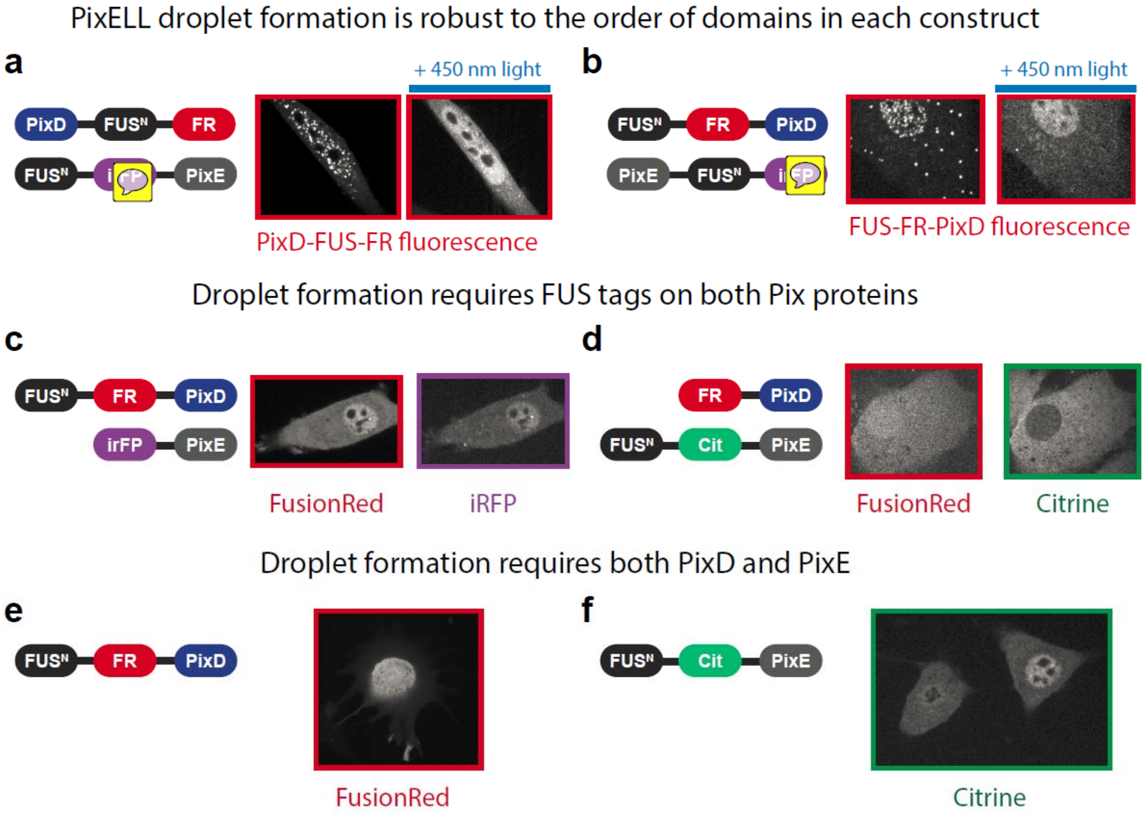


**Figure S2, related to Figure 2. Protein domain requirements for functional PixELL expression.** All panels show representative still images of cells expressing constructs indicated in cartoon diagrams. (**a,b**) PixD (**a**) and PixE (**b**) still achieve functional, light-switchable clustering when IDRs + fluorescent proteins are fused *via* C-terminal attachment. (**c,d**) Functional light-switchable clustering requires intrinsically disordered protein region (IDR) fusion to *both* PixE (**c**) and PixD (**d**). (**e,f**) Functional light-switchable clustering requires expression of both IDR-tagged PixD and PixE – cells expressing only one component exhibit diffuse localization.

#### Supplementary Figure 3

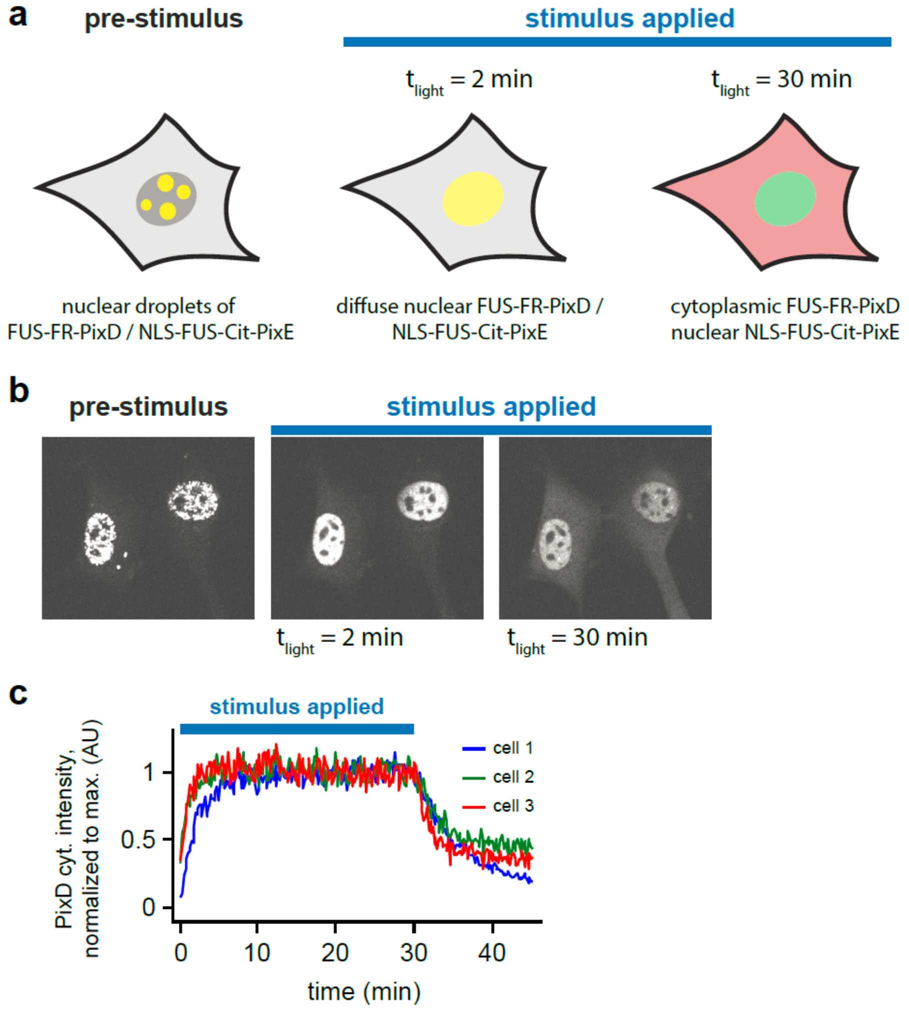


**Figure S3, related to Figure 3. Photoswitchable nuclear—cytoplasmic localization using subcellularly-localized Pix proteins.** (**a**) Schematic showing nuclear-localized PixELLs formed by $FUS^N$-FusionRed-PixD and NLS-$FUS^N$-Citrine-PixE. Droplets would be expected to dissolve shortly after light exposure, and PixD (which lacks an NLS) should be able to subsequently diffuse into the cytoplasm. (**b**) Experimental data of the constructs in **a** showing immediate dissolution of droplets, followed by redistribution of PixD into the cytoplasm. (**c**) Quantification of the cytoplasmic $FUS^N$-FusionRed-PixD intensity during the experiment shown in **b** for three independent cells.

#### Supplementary Figure 4

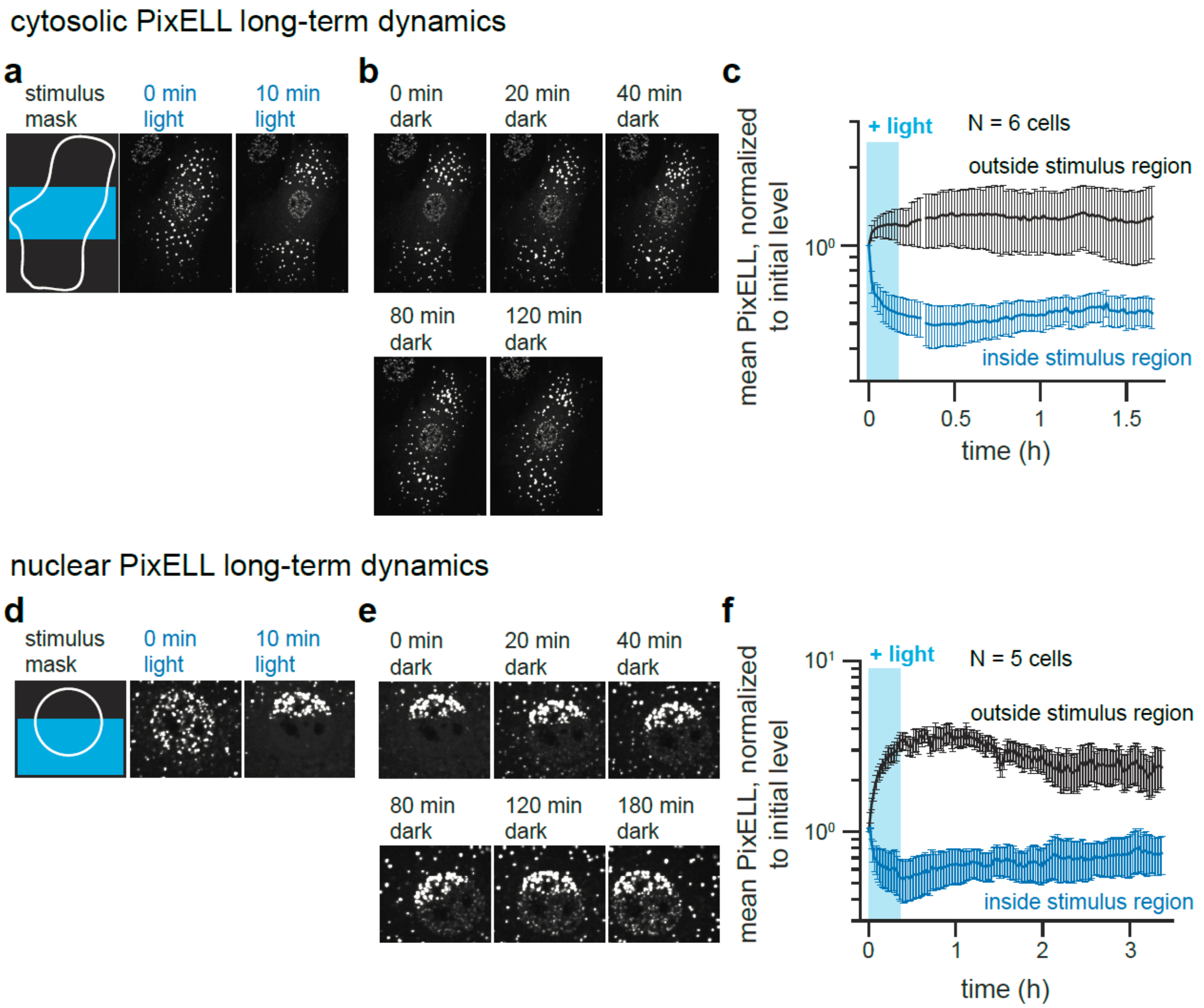


**Figure S4, related to Figure 3. Droplet-induced spatial asymmetry is long-lived.** (**a**) Schematic and images of local 450nm illumination in the cytosol of NIH 3T3 cells expressing the PixELL system. Fluorescent images of $FUS^N$-FR-PixD are shown both before and during light stimulation. (**b**) Images of the same NIH-3T3 cell are shown after transient illumination. (**c**) Quantification of Fold Change in cytoplasmic intensity inside and outside stimulation mask region. Mean + SEM for 6 cells are shown. (**d**) Schematic and images of local 450nm illumination in the cytosol of NIH 3T3 cells expressing the PixELL system. Fluorescent images of $FUS^N$-FR-PixD are shown both before and during light stimulation. (**e**) Images of the same NIH-3T3 cell are shown after transient illumination. (**f**) Quantification of Fold Change in cytoplasmic intensity inside and outside stimulation mask region. Mean + SEM for 5 cells are shown.

#### Supplementary Figure 5

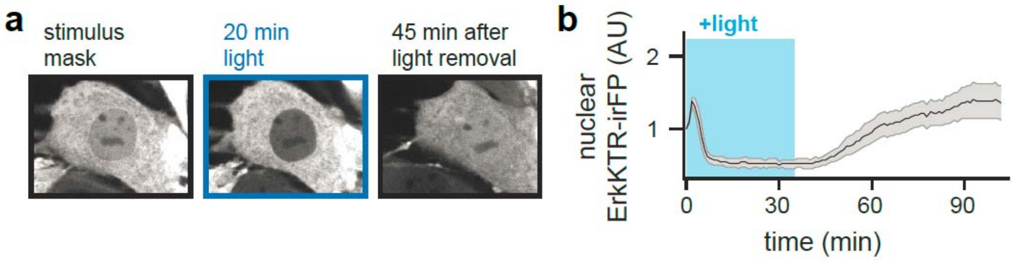


**Figure S5: FGFR1-optoDroplet control retraction in NIH-3T3 cells. Related to Figure 6. (a)** Images of Erk Kinase Translocation Reporter (KTR) during and after blue light stimulation of FGFR1-optoDroplet. **(b)** Quantification of nuclear ErkKTR via mean irFP fluorescence during and after blue light stimulation, for optoDroplet-FGFR1. Mean + SEM are shown for N=5 cells.

#### Supplementary Movie Legends

**Movie S1**
Kinetic Monte Carlo simulations showing aggregation for three different values of the interaction strength parameter θ. Simulations are each run for $10^5$ iterations (corresponding to approximately 18 simulated time units in each case). Three different values of θ are shown: $\theta = 0.2$ (solid-like), $\theta = 1$ (liquid-like) and $\theta = 2$ (diffuse). Related to Figure 1.

**Movie S2**
Kinetic Monte Carlo simulations illustrating the effect of stimulus-dependent local inhibition of aggregation (a local decrease in the value of θ). Simulation shows an initial aggregation phase (where $\theta = 1$ globally), followed by local disassembly ($\theta = 2$ on the right half of the lattice), followed by a return to global value of $\theta = 1$. Blue bar indicates the timing and spatial extent of stimulation. Related to Figure 1.

**Movie S3**
Long-term kinetic Monte Carlo simulations for $\theta = 1$. In the first section of the movie, clustering was initiated from diffuse initial conditions, leading to the growth of individual clusters by Ostwald ripening. In the second section, the simulation was initialized from a single droplet, demonstrating that no additional droplets form over long simulation times. Related to Figure 1.

**Movie S4**
Time lapse imaging of NIH3T3 cells expressing $FUS^N$-Citrine-PixE and $FUS^N$-FusionRed-PixD. Cells show repeated reversible formation and dissociation of clusters in blue light. Images were acquired with RFP imaging settings (561 nm excitation) every 30 sec for 5 h. Cells were stimulated with blue light from a digital micromirror device (DMD) for 10 min once each hour. Blue line in movie signifies timing of light delivery. Related to Figure 2.

**Movie S5**
Time lapse imaging of a NIH3T3 cell expressing $FUS^N$-Citrine-PixE and $FUS^N$-FusionRed-PixD, imaged in a subcellular region where clusters fuse over time. Images were acquired with RFP imaging settings (561 nm excitation) every 10 sec for 1 h. Yellow arrows indicate examples of fusion of distinct PixELL clusters. Related to Figure 2.

**Movie S6**
Time lapse imaging of an NIH3T3 cell expressing $FUS^N$-Citrine-PixE and $FUS^N$-FusionRed-PixD in response to transient, local blue light stimulation. Images were acquired with RFP imaging settings (561 nm excitation) every 30 sec for 1 h. Blue line in the video signifies both the timing of application and location of the blue light stimulus. Related to Figure 4.

**Movie S7**
Time lapse imaging of an NIH3T3 cell expressing of $FUS^N$-Citrine-PixE and $FUS^N$-FusionRed-PixD while cell is stimulated with a gradient of blue light. Images were acquired with RFP imaging settings (561 nm excitation) every 10 sec for 40 min. Cells were stimulated with a linear horizontal gradient of blue light intensity. Blue line in video signifies both the timing of application and orientation of the blue light gradient.

**Movie S8**
Time lapse imaging of approximately 10 µm x 20 µm patches of NIH-3T3 cell membranes expressing Myr-FUS$^N$-FusionRed-Cry2 (Myr-OptoDroplets). Cells were treated with DMSO as a control (part 1 of the movie), 1 µg/mL latrunculin A (part 2), or 0.5 µg/mL nocodazole (part 3). Continuous blue light is added at time zero. Scale bar indicates 10 µm. Membrane-localized droplets exhibit active transport in DMSO- and latrunculin-treated cells, with active transport dramatically reduced in nocodazole-treated cells. Related to Figure 5.

**Movie S9**
Time lapse imaging of a NIH-3T3 cell expressing MYR-Fus$^N$-FusionRed-Cry2 during local-to-global stimulation with blue light. Images were acquired with RFP imaging settings (561 nm excitation) every 30 sec for 1.5 h. Cells were first stimulated with a local 450 nm light input, followed by a switch to global illumination at the same intensity. The blue line in video signifies the timing and spatial range of blue light stimulation. Related to Figure 5.

**Movie S10**
Time lapse imaging of a NIH-3T3 cell expressing MYR-Fus$^N$-FusionRed-Cry2-FGFR1 during alternating bouts of global stimulation with blue light and withdrawal of the blue light stimulus. Images were acquired with RFP imaging settings (561 nm excitation) every 30 sec for 1 h. The blue line in video signifies the timing of blue light stimulation. Related to Figure 6.

**Movie S11**
Time lapse imaging of a NIH-3T3 cell expressing MYR-Fus$^N$-FusionRed-Cry2-FGFR1 during local stimulation with blue light. Images were acquired with RFP imaging settings (561 nm excitation) every 30 sec for 54min. Cells were first stimulated with a local 450 nm light input in two different locations (on the two horizontal sides of the cell). The blue lines in the video signifies the timing and spatial range of blue light stimulation. Related to Figure 6.

**Movie S12**
Time lapse imaging of a NIH-3T3 cell expressing MYR-Fus$^N$-FusionRed-Cry2-FGFR1 during local-to-global stimulation with blue light. Images were acquired with RFP imaging settings (561 nm excitation) every 30 sec for 1h. Cells were first stimulated with a local 450 nm light input, followed by a switch to global illumination at the same intensity. The blue line in the video signifies the timing and spatial range of blue light stimulation. Related to Figure 6.

#### References


Kim, N., Kim, J.M., Lee, M., Kim, C.Y., Chang, K.Y., and Heo, W.D. (2014). Spatiotemporal control of fibroblast growth factor receptor signals by blue light. Chemistry & biology *21*, 903-912.

Regot, S., Hughey, J.J., Bajar, B.T., Carrasco, S., and Covert, M.W. (2014). High-sensitivity measurements of multiple kinase activities in live single cells. Cell *157*, 1724-1734.

Serebrinsky, S.A. (2011). Physical time scale in kinetic Monte Carlo simulations of continuous-time Markov chains. Physical review E, Statistical, nonlinear, and soft matter physics *83*, 037701.

Shin, Y., Berry, J., Pannucci, N., Haataja, M.P., Toettcher, J.E., and Brangwynne, C.P. (2017). Spatiotemporal Control of Intracellular Phase Transitions Using Light-Activated optoDroplets. Cell *168*, 159-171 e114.

Toettcher, J.E., Gong, D., Lim, W.A., and Weiner, O.D. (2011). Light-based feedback for controlling intracellular signaling dynamics. Nature methods *8*, 837-839.

Toettcher, J.E., Weiner, O.D., and Lim, W.A. (2013). Using optogenetics to interrogate the dynamic control of signal transmission by the Ras/Erk module. Cell *155*, 1422-1434.
